# Laser NIR irradiation enhances antimicrobial photodynamic inactivation of biofilms of *Staphylococcus aureus*

**DOI:** 10.1101/2024.05.08.593196

**Authors:** Leandro Mamone, Roberto Tomás, Gabriela Di Venosa, Lautaro Gándara, Edgardo Durantini, Fernanda Buzzola, Adriana Casas

**Author notes:** CorrespondingAuthor: Universidad de Buenos Aires, CONICET, Hospital de Clínicas José de San Martín,Centro de Investigaciones sobre Porfirinas y Porfirias (CIPYP), Ciudad de Buenos Aires, Argentina.

## Abstract

Photodynamic inactivation (PDI) utilizes a photosensitizer (PS) activated by visible light to generate reactive oxygen species (ROS), to kill bacteria. PDI is effective against planktonic microorganisms, but biofilms are less sensitive due to limited PS and oxygen penetration. Near-infrared treatment (NIRT), involve the use of near-infrared light to kill bacteria either via thermal effects or ROS production.

Our objective was to enhance *S. aureus* biofilm’s sensitivity to PDI by pre-treating with NIR irradiation before visible light exposure.

In an *in vitro* biofilm model, laser NIRT (980 nm) followed by exposure to PDI, showed a synergistic effect on bacterial viability loss (4-log CFU vs 1-log loss with individual treatments). Interestingly, pre-heating liquid medium had no significant impact on PDI efficacy, suggesting that both thermal and non-thermal effects of NIR may be involved.

NIRT increased PS uptake, induced clefts in the biofilm matrix, and released bacterial cells from the biofilm. NIRT induced a transient increase in the temperature to 46°C of *in vitro* cultures, however under the same conditions, when mice were irradiated, skin temperature rose to 37°.

Our findings suggest that NIR irradiation serves as a complementary treatment to PDI, allowing reducing PS concentration, and highlighting its potential as an effective and resource-efficient antibacterial approach.

## Introduction

Photodynamic Inactivation (PDI) is a powerful technique used to eliminate microorganisms through the combined action of a light-activated drug called a photosensitizer (PS), visible light, and molecular oxygen leading to the formation of reactive oxygen species (ROS). These ROS can induce oxidative damage to lipids, proteins, and nucleic acids, thereby effectively eliminating microbial cells, viruses, and parasites [1,2].

The application of PDI against bacteria has shown promising results in the treatment of various superficial infections and bacteria-caused diseases. These include periodontal and endodontic infections, acne, non-healing ulcers, chronic skin wound infections, chronic rhinosinusitis, otitis media, prosthetic joint infections, and the potential treatment of colonised catheters and other medical devices [1,3,4].

In addition to medical applications, PDI has also been utilized in wastewater treatment, disinfection of food products, and even in the treatment of infectious microorganisms associated with blood transfusions and ventilator-associated pneumonia [3,5–8].

Due to its wide range of biochemical targets, PDI can be used to treat all types of microbial infections, including those caused by antibiotic-resistant strains [9–11].Interestingly, PDI-tolerant *Staphylococcus aureus* exhibited increased susceptibility to antibiotics [12].

A wide range of substituted cationic porphyrinshave been used to mediate PDI of diverse species of pathogens [13–15]. Ithas also been reported that cationic PSs are particularly efficient at eradicating biofilms developed by Gram-positive bacteria[9].

Our group has previously described that 5,10,15,20-tetrakis[4-(3-*N*,*N*-dimethylaminopropoxy)phenyl]porphyrin (TAPP) can eradicate planktonic cultures of *S. aureus* upon irradiation with visible light. However, under the same conditions, biofilm viability was only reduced in 1-log [16].

Biofilm-associated pathogens account for up to 80% of all microbial infections in humans and are up to 1000-fold tolerant to antimicrobial treatments due to multiple factors including the reduced penetration of antimicrobials through the matrix, which thus represents a diffusion barrier that protects bacterial cells [17]. PDI is capable of eliminating or inducing substantial reductions in the microbial community, of both planktonic and biofilms-forming bacteria[16,18]. Nevertheless, biofilms are less sensitive to PDI than planktonic bacteriamainly due to the reduced penetration of the PS and oxygen into the biofilm and bacterial cells [7,19].

Various biochemical methods have been proposed to tackle the challenge of disrupting the architecture of biofilms, including matrix-targeting enzymes, bacteriophage therapy, quorum-sensing inhibitors, as well as small molecules[20].

While these approaches have shown promise, a physical method that interferes with biofilm integrity presents certain advantages. Physical forces tend to be independent of bacterial and matrix composition, which makes them more universally applicable [21,22]. In this regard, the disruption of biofilms using techniques such as ultrasound, shock waves, or thermal treatments has been reported, with a particular focus on the elimination of *S. aureus* biofilms [21–26].These physical methods offer a potential solution for combating and eliminating biofilms in different environments.

In recent years, near-infrared treatments (NIRT) have emerged as promising antibacterial tools, particularly in combating resistant strains. These treatments utilize near-infrared (NIR) radiation within the range of 700-1300 nm, either alone or in combination with NIR-sensitive materials, to generate heat and ROS[27–29].

NIRT can cause damage to bacteria throughthe absorption of photons by water, which increases the vibrational energy and raises the temperature of bacterial cells, leading to their damage (thermal effect). Alternatively, the absorption ofphotons by specific chromophores, either inside or outside of the bacteria, leads to the production of reactive oxygen species (ROS) which damage cellular components such as membranes or biofilm matrix. This is referred to as the non-thermal effect[30,31].The thermal effects can be replicated by simply warming the irradiated sample to the temperature that would have been reached by the NIR exposure. Since light-based therapies are non-invasive, selective and highly effective, PDI and NIRT have shown great potential to combat bacteria embedded in biofilms [32].

We have previously reported that both NIRT with a 980-nm laser and PDI employing toluidine blue and a 635-nm laser individually resulted in 1-log reduction in viable counts of *S. aureus* RN6390 biofilms. The consecutive application of 980-nm and 635-nm laser treatments on TB-treated biofilms exhibited additive but not synergistic effects[33]. In the present work, we aimed to establish conditions under which biofilm dispersal occurs atlower temperatures, employing other PS, and toachieve synergistic effects when combined with PDI.

Our goalwas to pre-treat the biofilms with NIR irradiation before visible light irradiation, to restore the sensitivity of bacterial cells to TAPP-PDI. This combination approach aims to achieve a biofilm-disrupting effect and improve the overall efficacy of TAPP-PDI against *S. aureus* biofilms.

## Results

### Determination of NIR irradiation parameters to treat biofilms

To set the conditions to be employed in the combined treatment with PDI, we tested NIR irradiation effects over biofilm cultures of *S. aureus* RN6390 (Figure 1A). Since water can absorb NIR energy, we first measured the temperature rise of Phosphate-buffered saline (PBS) during and after 30 seconds of 980 nm laser exposure at different output powers in continuous mode. The temperature (initially at 24.5 °C) increased in a dose-response manner during the irradiation (grey-shaded region) and decreased gradually after treatment. At 2.5 W/cm^2^ the temperature never exceeded 35 °C (ΔT = 10 °C) and when the radiance was set at 5 W/cm^2^, the ΔT was 17.2 °C (maximum of 41.7 °C). In addition, at the highest light dose employed (7.5 W/cm^2^), the temperature reached 37 °C after 15 seconds of irradiation, and the maximum temperature attained was 46.0 °C, ΔT = 21.5 °C (Figure 1. A).

**Figure 1.**
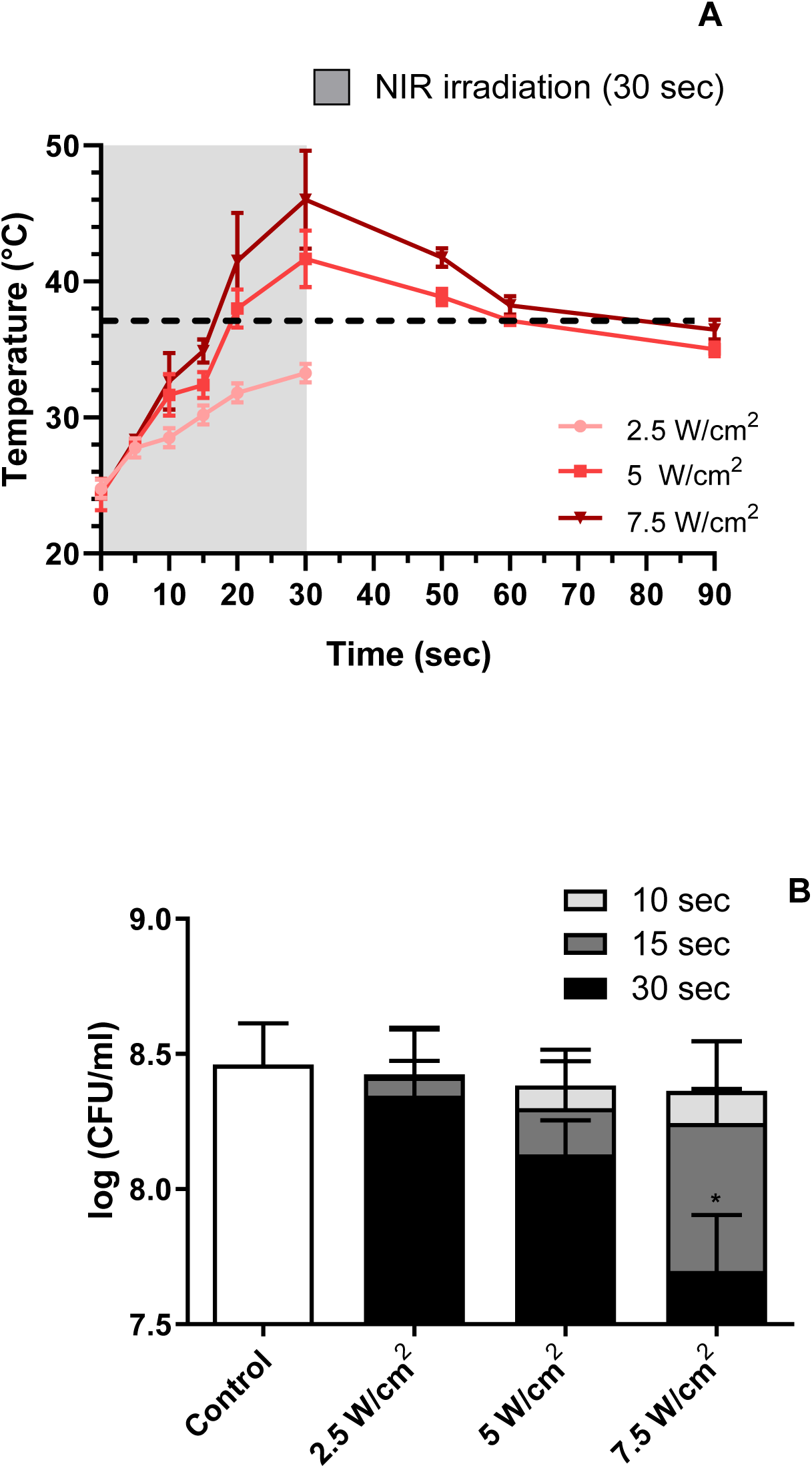
Temperature rise induced by NIRT and its effect on *S. aureus RN6390* biofilm cultures. **A)** Temperature of PBS over biofilm cultures during NIR irradiation. A 980nm laser was employed at different output powers for 30 sec. on continuous mode and temperature registers were obtained during and after irradiation. Gray shaded area: laser emission on. Dotted line: 37 °C. **B)** Bacterial viability after NIRT employing different powers and exposure times. Data are means of 4-6 replicates. Control: non-treated bacteria. *p<0.01 ANOVA, followed by Dunnet’sposthoc test as compared to the control.

Subsequently, we investigated the impact of NIRT on biofilm viability, as depicted in Figure 1B. Noteworthy results emerged, indicating a significant reduction in bacterial viability exclusively under the most extreme conditions—7.5 mW/cm² for 30 seconds—when compared to the untreated control. Remarkably, employing these parameters, the viability of the NIR-treated biofilm diminished to approximately 20% of the untreated control (p = 0.01).

### Combination of NIRT and PDI to reduce biofilm viability

To enhance the efficacy of PDI on biofilm viability, utilizing TAPP as the photosensitizer, we combined NIRT with TAPP-PDI in the experimental framework illustrated in Figure 2A. Briefly, TAPP was introduced into the medium, and the biofilms underwent treatment with NIRT, employing the conditions previously examined in Figure 1. Subsequently, a 1-hour dark incubation period ensued, followed by subsequent irradiation with visible light.

**Figure 2.**
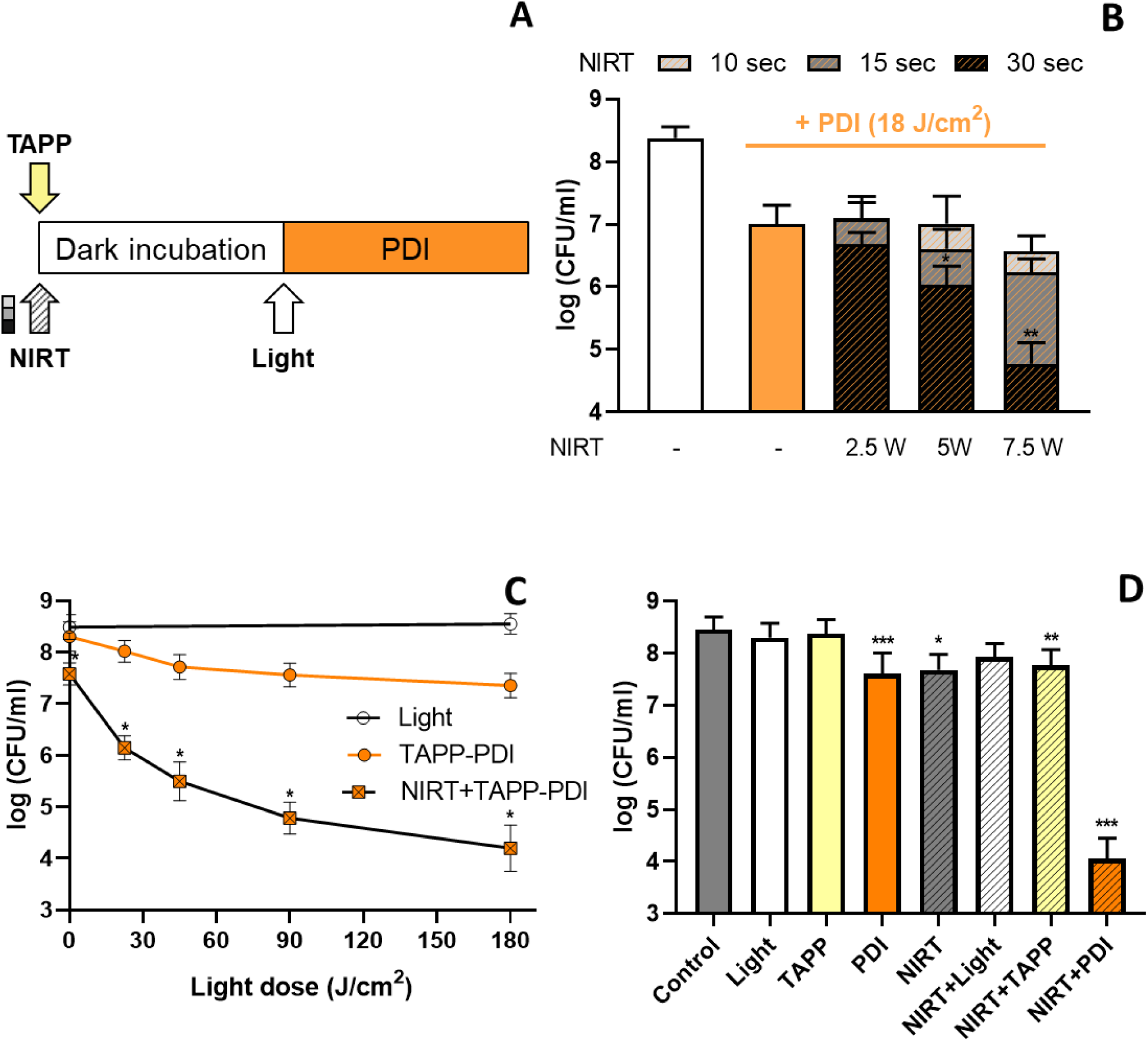
Combination of NIRT and TAPP-PDI on the viability of *S. aureus* RN6390biofilms. **A)**Experimental design. NIRT: 2.5, 5 or 7.5 W/cm^2^, continuous mode. Light: visible light (180 J/cm^2^), 2.5 µM TAPP, 1 hour. **B)** Bacterial viability after TAPP-PDI alone or combined with different doses of NIRT.**C)** Effect of the light dose of PDI on the combination with NIRT (30 sec, 7.5 W/cm^2^). **D)** Viability of biofilms treated with NIRT (30 sec, 7.5 W/cm^2^) followed by TAPP-PDI and controls of visible light and TAPP previously exposed to NIRT.Data are means of 4 to 6 biological replicates. Two-way ANOVA with Tukey posthoc test for B and C as compared to TAPP-PDI treatment, and one-way ANOVA with Tukey posthoc test as compared to the non-NIRT treatments for D.

The conditions selected to carry out TAPP-PDI in the following experiments were 2.5 µM TAPP and 180 J/cm^2^ of visible light. As we previously reported [16], under these conditions, TAPP-PDI induced a 1-log reduction in biofilm viability.

However, upon combining TAPP-PDI with NIRT, the viability reduction depended on the NIRT irradiance conditions and exposure time (Figure 2B). When PDI and 2.5 W/cm^2^ NIRT were employed (regardless of the exposure time), no significant differences were obtained, as compared to PDI alone. On the other hand, when PDI was combined with NIRT at irradiances of 5 and 7.5 W/cm^2^ during 30 sec, a significant reduction in biofilm viability was attained (2-logs for 5 W/cm^2^, and 4-logs for 7.5 W/cm^2^ respect to the untreated controls). The statistics of the comparison among biofilms treated with NIRT, PDI, and both therapies indicate that the combination of NIRT and PDI results in a synergistic effect rather than an additive one (two-way ANOVA test) when using power levels of 7.5 W/cm^2^ for 30 sec as well as when employing 5 mW for 30 sec (p < 0.0001 and p=0.03 respectively) (Figure 2C).Moreover, previous exposure of biofilms to NIRT synergizes the antibacterial action of PDI in all the 15 to 180 J/cm^2^ range (Figure 3C).Controls of the different treatments are shown in Figure 2D.

**Figure 3:**
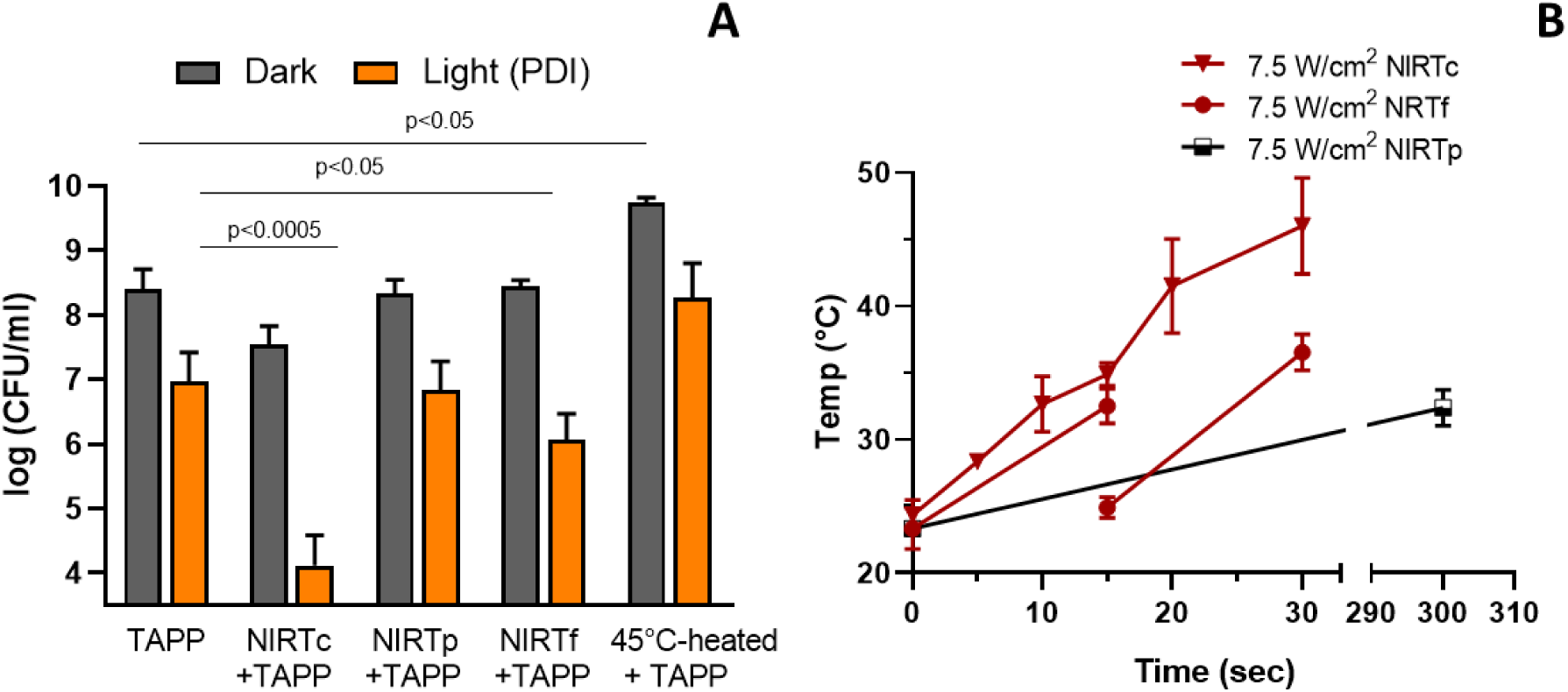
Viability of *S. aureus* biofilms treated with 225 J/cm²NIRT under different schemes. **(A)**Biofilms were exposed to TAPP and immediately afterwards NIRT was applied in continuous mode (30 sec, 7.5 W/cm^2^), pulsed mode (7.5 W/cm^2^, 2 Hz, 50 msec pulse width, 300-sec irradiation)or continuous mode fractionated, (15 sec ⋏+ 120-sec recovery + 15 sec ⋏, 7.5 W/cm^2^) with or without ulterior PDI (180 J/cm^2^).Additionally, biofilms were exposed to PBS containing TAPP heated to 45°C. **(B)**Temperature increase of PBS as a function of the time exposure to different NIRT schemes.

### Influence of temperature on the enhancing NIRT effect on PDI

In exploring the influence of temperature on theenhancing NIRT effect on PDI, we fine-tuned the NIRT conditions to deliver an energy dose equivalent to that applied in the 7.5 J/cm² over 30 seconds scenario, that is a dose of 225 J/cm² of NIRT on continuous mode (NIRTc) already employed in Figure 2,while ensuring the attainment of the lowest possible maximum temperature (Figure 3).To facilitate the dissipation of energy delivered by the NIR laser into the medium we employed two approaches: NIRT in pulsed mode (NIRTp) and fractionated doses of continuous NIRT(NIRTf). For the pulsed mode scheme, we did not observe any significant impact on bacterial viability or enhancement of TAPP-PDI. The maximum temperature recorded under these conditions was 32.4°C (Figure 3B).Subsequently, we divided the exposure time into two intervals of 15-sec each, separated by a 120-sec interval. Despite these conditions, we did not observe any notable impact of the fractionatedscheme on bacterial viability. However, a significant effect was observed in enhancing TAPP-PDI, although the potentiation effect was less pronounced thanin the non-fractionated treatment (2 logs versus 4 logs). The maximum temperature recorded under these conditions was 36.5°C.

To further explore the impact of the maximum temperature reached by the medium during NIRT, we introduced PBS (supplemented with TAPP) previously heated to 46°C to biofilm cultures andincubated them in the dark at 37°C for 1 hour. Surprisingly, no significant enhancement of TAPP-PDI was observed. Furthermore, biofilms treated with heated-TAPP and not exposed to visible light exhibited a significant increase in CFU/ml count compared to the non-heated control.

### *In vivo* irradiation of mice skin under the NIRTcsetup conditions

We conducted an experiment using the NIRTc setup conditions to irradiate shaved skin from previously anaesthetized CF-1 mice. The initial basal temperature on the mice’s skin measured 36°C, which dropped to 31°C after anaesthesia. Subsequently, we employed a cooling spray as an additional method to further reduce the temperature before NIRT, minimizing the potential adverse effects of irradiation. This additional cooling step brought the temperature down to 20°C. The temperature recording on the irradiated skin, as the NIRT procedure advanced, is depicted in Figure 4. NIRT progressively elevated the skin temperature during irradiation, reaching 37 °C (ΔT=17°C) at 30 seconds of irradiation. Importantly, no harmful effects were observed on the skin immediately after irradiation or in the subsequent days.

**Figure 4.**
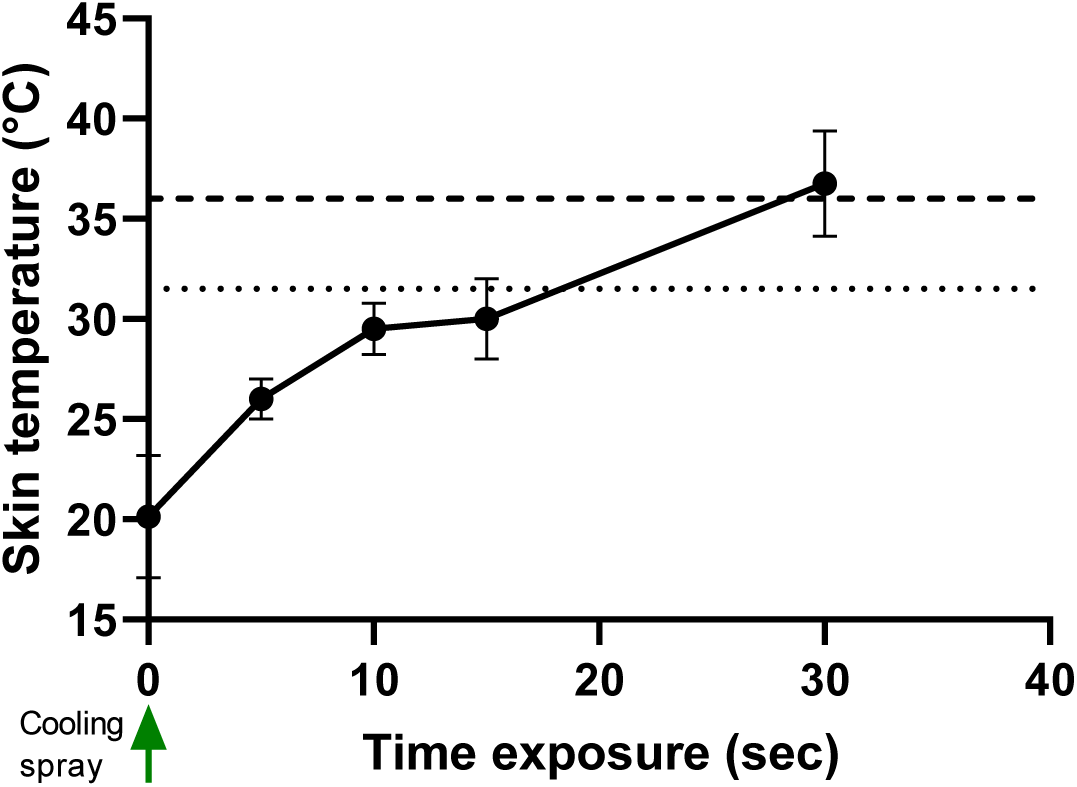
Mice skin temperature during *in vivo*NIRTc. CF-1 mice were irradiated with the 980 nm laser at 7.5 W/cm^2^ for 30 seconds, after being anesthetized. A cooling spray was used to reduce the temperature in the irradiated skin region before irradiation Dashed line: basal skin temperature before anaesthesia. Dotted line: skin temperature after anaesthesia. Values are means ± SD of n=3 mice.

Therefore, considering the reduction of viability attained, we selected NIRT on continuous mode at 7.5 W/cm^2^ for 30 sec., as the conditions to perform further experiments, because those were the conditions under which NIRT showed the greatest enhancing effect on TAPP-PDI, keeping the temperature below the temperatures generally accepted in clinical treatments. Therefore, unless stated, we will refer to NIRT as the continuous mode of irradiation.

### NIRT effects on biofilms and PDI enhancement

#### TAPP uptake into biofilms

To explore the underlying causes of the enhanced TAPP-PDI effect by NIRT, we measured the overall incorporation of TAPP into the biofilm after NIRT. Since the ratio of TAPP per bacteria in NIRT-treated biofilms (22.6 nmoles TAPP/10^8^ cells) increased significantly compared to the non-NIRT treated ones (17.4 nmoles TAPP/10^8^ cells)we conclude that NIRT is capable of increasing the uptake of TAPP by the biofilms (data not depicted).

#### NIRT-induced detachment of bacterial cells from biofilms

To further determine whether NIRT was capable of detaching bacterial cells from attached biomass, the air-liquid interface (ALI) was removed from biofilm cultures and both phases (attached biofilm and non-attached/planktonic bacteria or ALI) were treated with NIRT and TAPP-PDI (Figure 5).While in the untreated biofilm cultures, the subpopulation of attached bacteria was almost doublethat of the bacteria subpopulation in the ALI, this ratio was reversed after NIRT, making bacteria in the ALI the most abundant subpopulation. It is worth mentioning that the total number of viable bacteria in the entire culture, as already mentioned, decreased after NIRT.On the other hand, when measuring the effects of NIRT on the attached biofilm biomass through crystal violet staining, a significant 12% reduction was detected compared to the untreated control (data not depicted).

**Figure 5.**
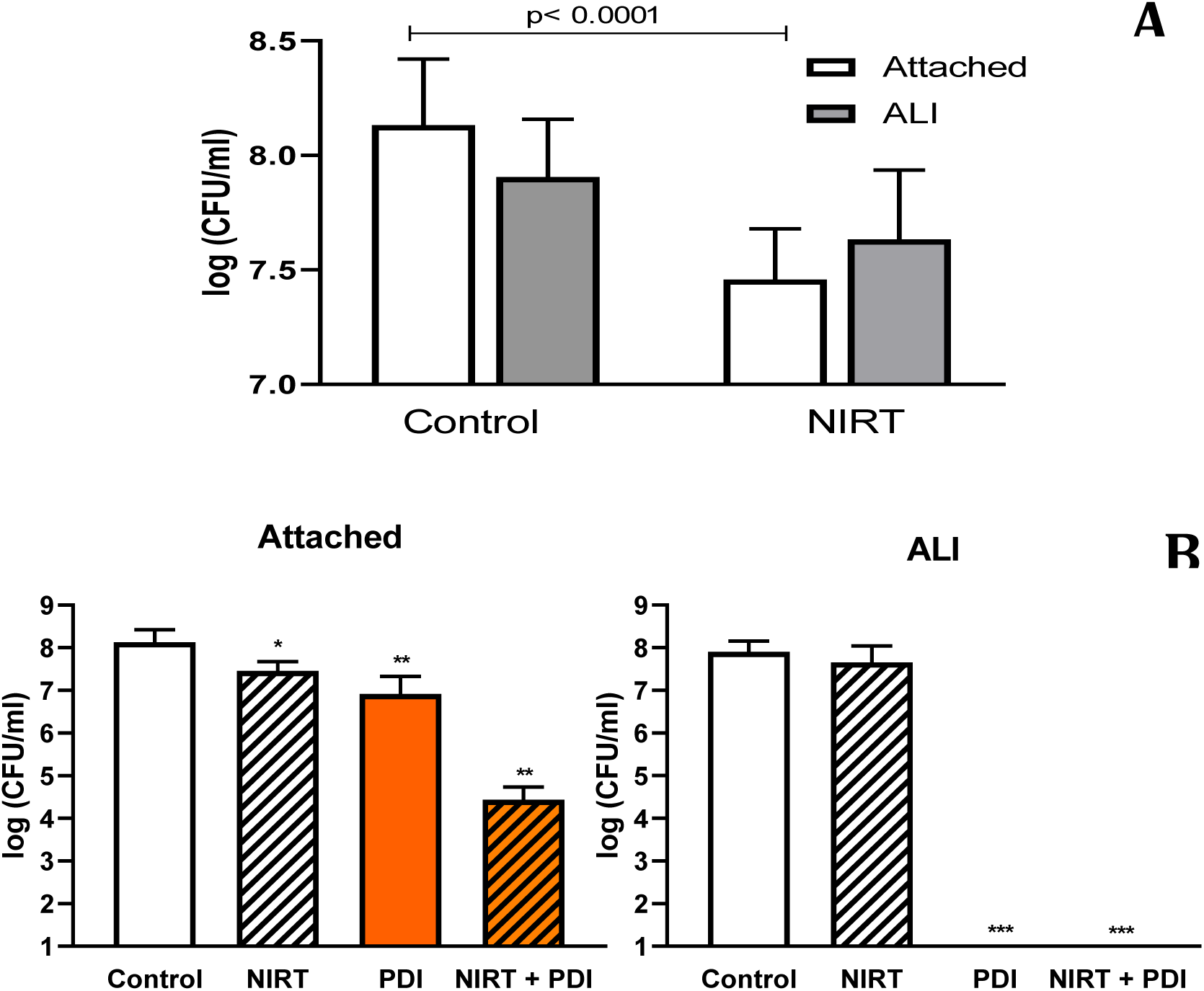
NIRT and TAPP-PDI effect on the viability of adherent or ALI biofilms of *S. aureus*. **A)** Viability of attached bacteria and ALI subpopulations after NIRT. **B)** Viability after combined treatments for each subpopulation. Data are means of 4 to 6 biological replicates. ANOVA, Dunnet post-hoc test as compared to the controls.

#### Effects of combined NIRT and TAPP-PDI treatment on different subpopulations of bacterial cells within biofilms

Following this, we conducted TAPP-PDI on the adhered biofilm phase, resulting in a 1-log reduction in bacterial viability relative to the untreated control. Notably, the combination of NIRT and TAPP-PDI yielded a more substantial impact, impairing viability by 4 logs. Furthermore, TAPP-PDI exhibited complete eradication of ALI bacteria, irrespective of prior NIRT exposure, as illustrated in Figure 5B.

#### SEM images of the NIRT and PDI-treated biofilms

SEM image*s of S. aureus* biofilms after TAPP-PDI correlated well with colony count data, and revealed that the treatmentwas capable of detaching adhered bacteria from the substrate (Figure 6). In addition, in TAPP-PDI-NIRT treated biofilms,massive bacteria detachment higher than the effect elicited by both treatments separately was observed after the combined treatment, in agreement with the CFU/ml counts. NIRT induced disruption across the biofilm matrix, which can be observed as notches in the biofilm structure (see inset).

**Figure 6.**
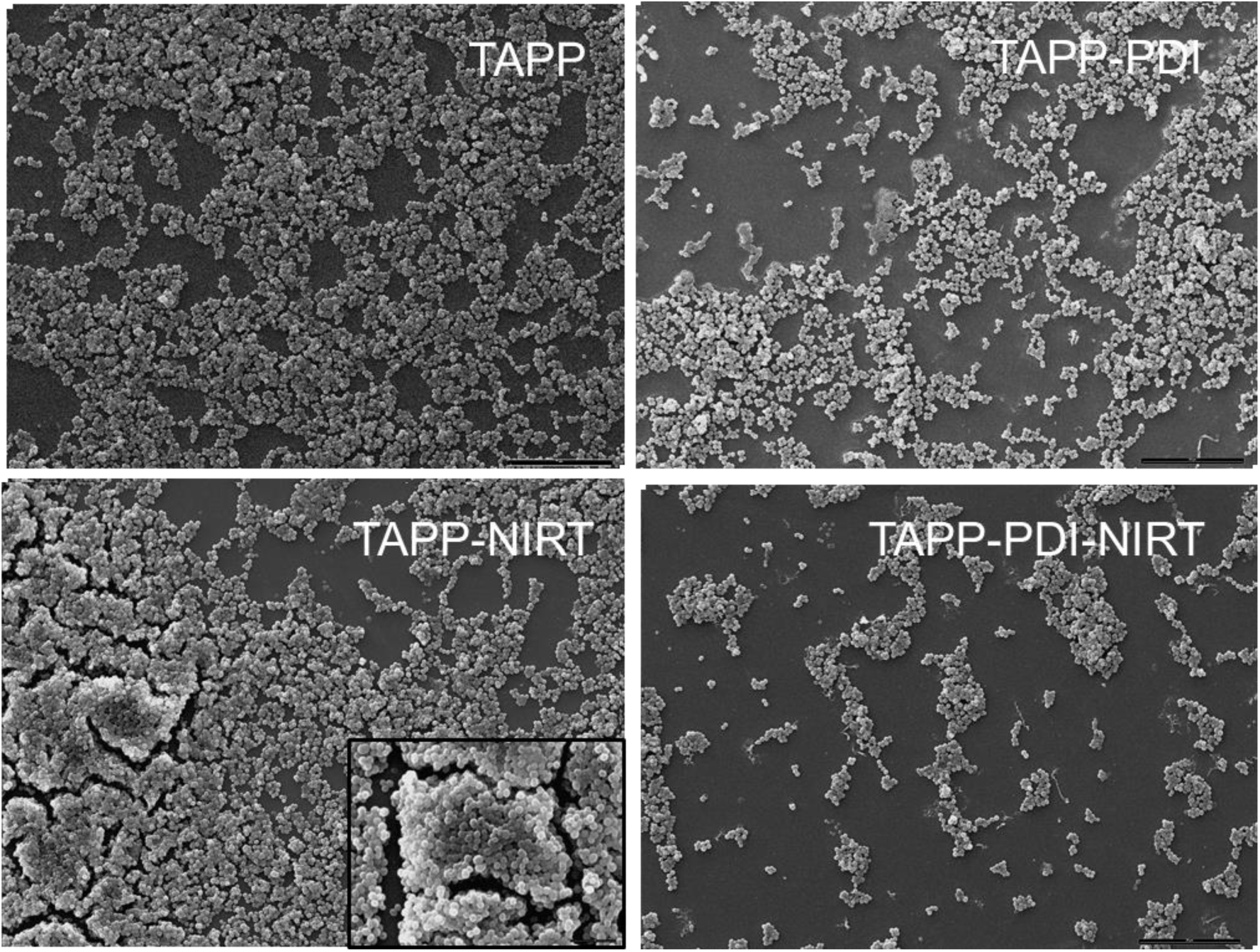
SEM images of biofilms treated with NIRT and TAPP-PDI. All the biofilms were treated with TAPP (2.5 µM) and also received either NIRT (225 J/cm^2^), PDI (180 J/cm^2^) or a combination of PDI and NIRT. Samples were fixed immediately after treatment. Magnification, 1000x. A representative image of each treatment out of 3 is depicted. The inset of TAPP-NIRT shows an image of 4000X magnification.

#### Analysis of NIRT and PDI effects on bacterial viability by CLSM

We also evaluated the viability of treated biofilms by CLSM using the LIVE/DEAD BacLight Bacterial Viability Kit (Figure 7). CLSM images of TAPP-incubated biofilms showed multilayered clumps of bacteria where the majority of cells were viable (green fluorescence). In contrast, biofilms treated with TAPP-PDI and/or NIRT showed a higher proportion of dead red fluorescent cells and a thinner layer of biofilm biomass. The characteristic feature of both treatments exposed to NIRT is the presence of clusters of greater size than TAPP-incubated control and TAPP-PDI-treated biofilms.

**Figure 7.**
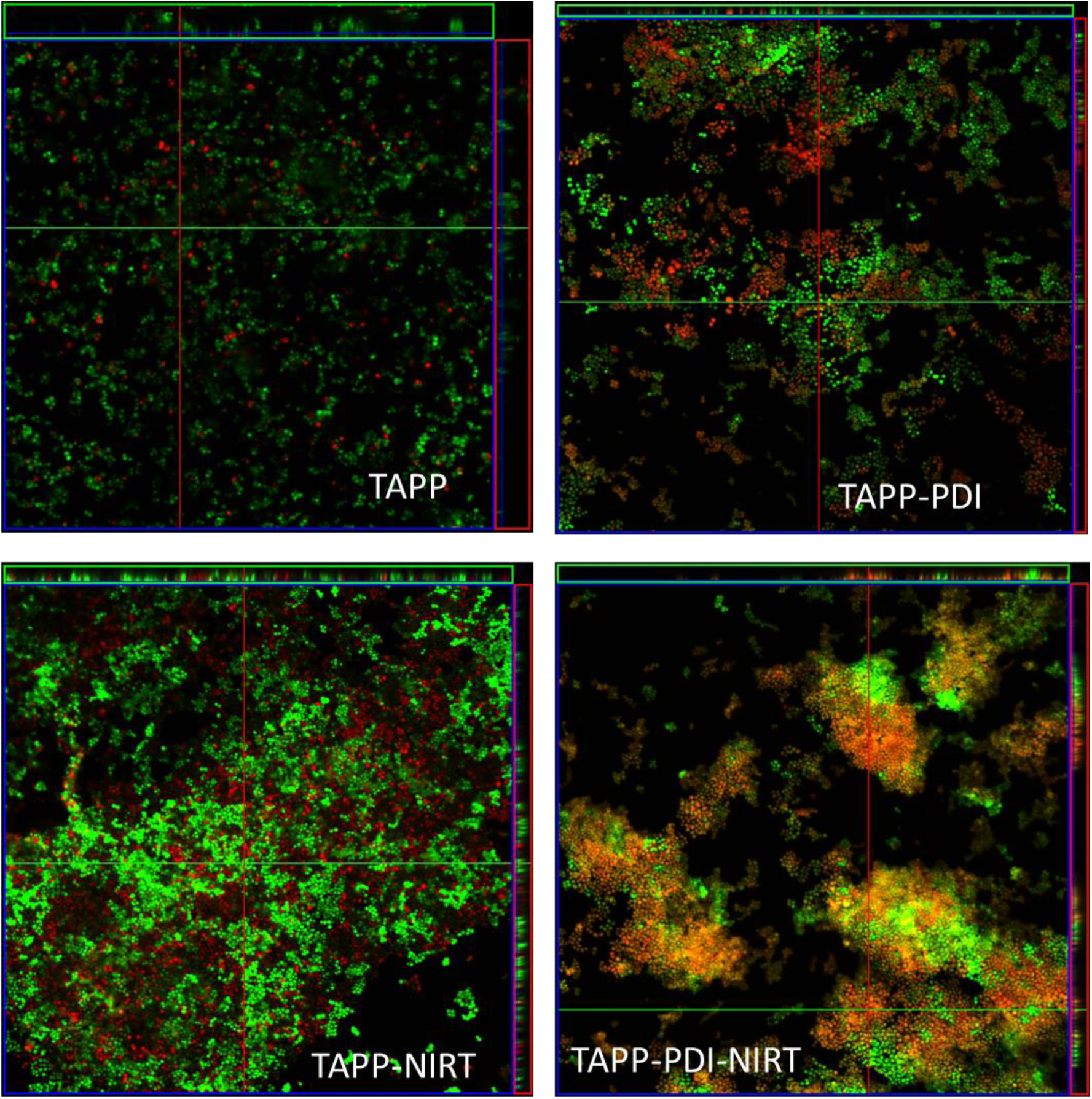
CLSM imaging analysis of TAPP-PDI and NIRT combined treatment on *S. aureus* biofilms. Viability after treatments was determined by IP/SYTO9 staining and confocal microscopy. All the biofilms were treated with TAPP (2.5 µM) and also received either NIRT (225 J/cm^2^), PDI (180 J/cm^2^) or a combination of TAPP-PDI and NIRT. A representative image of each treatment out of 3 is depicted.

## Discussion

The biofilm structure serves as a barrier to antimicrobial treatments. Consequently, a therapeutic approach capable of disrupting the integrity of the biofilm, facilitating improved drug penetration, would constitute a noteworthy advancement with promising clinical applications [7,21].

Currently, the gold standard for removing biofilm from chronic wounds is repeated sharp debridement, an often painful and imprecise method. To address this, mechanical removal is combined with antibiotics. However, these recalcitrant wounds often do not heal, due to ineffective debridement and emergence of antibiotic resistance [36]. Thus, a laser/light-based treatment to locally disrupt biofilm integrity and kill biofilm-associated bacteria could represent a promising alternative.

Particularly, NIRT could be useful in the treatment of slow-growing or persistent bacteria, reversing the conditions that promote the generation of cells with low metabolism, and therefore, making them sensitive to PDI or other antimicrobial treatments. It would also allow for overcoming the diffusion barrier that the EPS matrix represents for most antimicrobial agents. It should be noted that the bacteria that NIRT could release from the extracellular matrix in infected tissues would also be exposed to the action of the immune system or circulating antibiotics.

Conversely, NIR radiation, with the ability to penetrate tissues up to 3 cm, is generally considered non-harmful [37]. However, a drawback of the technique is the shedding of viable bacteria—capable of infecting new tissues—that NIRT cannot effectively inactivate. In addressing this limitation, PDI emerges as a valuable tool capable of efficiently inactivating various bacterial species, including those resistant to clinically used antibiotics. This positions PDI as a valuable complement to a combined treatment involving NIR laser irradiation for clinical infections.

In this work, we used the porphyrin TAPP as the PS for the elimination of *S. aureus* biofilms, an important human pathogen that causes a wide range of clinical infections. The tetrapyrrole macrocycle of TAPP is substituted by four (dimethylamino)propoxy)phenyl groups at the *meso* positions. Terminal amino substituents can acquire positive charges by protonation in aqueous medium, increasing the interaction with the cell envelope of microorganisms [38]. This photosensitizer was effective in inactivating bacteria and yeast in cell suspensions [16,39,40]. Herein, we were able to reverse the initial treatment resistance of *S. aureus* biofilms, by combining TAPP-PDI with NIR laser irradiation.

Noteworthy, the synergism between NIRT and PDI resulted in 4-log reduction in the viability of biofilm cultures. This achievement stands out compared to our prior study [16], where a 2.5 μM TAPP concentration only led to a 1-log reduction. In that earlier investigation, it was imperative to elevate the TAPP concentration to 20 μM to attain a 2-log reduction in the viability of biofilm cultures grown under comparable conditions.

Any novel method must demonstrate a 3-log reduction in CFU efficacy before being designated as bactericidal [9]. In this study, we achieved a significant advancement by successfully attaining a 4-log reduction through the combined application of NIRT and PDI, a milestone that remained unattainable through the use of separate treatments.

Enhancement of antimicrobial blue light treatment using endogenous PSs was previously achieved upon treatment of S*. aureus* biofilms with NIR with spectral range (780-1060nm) [41]. However, there are no previous reports on the combination of PDI employing exogenous PSs and NIRT.

The observed synergy among our radiation-based treatments in the experimental setup was somewhat linked to the maximum temperature attained during NIRT. This effect was triggered by the delivery of 227J/cm^2^ in continuous mode, whereas pulsed or fractionated schemes failed to induce the same outcome.However, it did not exclusively hinge on temperature, as evidenced by the absence of synergistic effects when heating the bacterial culture medium (without irradiation) to the maximum temperature achieved during NIRT, followed by subsequent PDI. Non-temperature-dependent effects triggered by the absorption of NIR by cellular compounds, such as cytochromes or certain protein complexes may also be acting [42]. However, the contribution of heat and photochemical processes involved in NIRT is still a matter of discussion. It was reported that upon irradiation of suspensions of *E. coli,* no significant differences were found in the killing rates between 940 nm laser treatment and water-based heating. Therefore, the authors concluded that the most important parameter involved in bacteria-killing is the maximum temperature achieved [37]. On the other hand, the bactericidal action of Nd:YAG laser light at 50 °C waspartly ascribed to thermal heating and partly to an additional, as yet undefined, mechanism [43].

Regarding the light scheme, in contrast to our observations, other authors found that with the application of fractionated or pulsed doses of 980 nm laser, a reduction of almost 3 logs in multispecies biofilms grown on titanium surfaces was attained, independently of the light dose scheme [44].

NIRT-induced photothermal therapy typically demands temperatures exceeding 45 °C for efficient bacterial eradication [45,46]. Structural changes generated by temperature promote gene expression to overcome cellular stress through a heat shock response that attempts to mitigate the effects of high temperatures until eventually the damage to the bacterial cell becomes irreversible [47].The therapeutic window for NIR diode lasers—where bacterial cell death is achieved without harming the host’s tissue—is quite limited. Maintaining the laser energy at a minimum is essential, as the temperature in the irradiated area increases rapidly[48].Tissue damage is related not only to the original temperature rise but also to the period for which the tissue is subjected to the increased temperature. Besides, the effects of NIR irradiation on tissue temperature increase depend on the thermal properties of the tissue, such as thermal conductivity, heat capacity, and local blood perfusion [49].

In our study, we observed a temperature increase of 17°C on the mouse skin surface when subjected to a continuous-mode 980 nm laser at 270 J/cm², peaking at 37°C. In human skin, a 100 ms spurt with cryogen spray resulted in a surface temperature reduction of 31°C and a cooling of 12°C at a 100 µm depth [50]. Consequently, there is a high likelihood of replicating our NIRT conditions on human skin without inducing hyperthermia, provided a cooling agent is applied beforehand.

Mild hyperthermia has been proven to induce the detachment of *in vitro* biofilms of *S. aureus,S. epidermidis*, and *K. pneumoniae.* It has been reported to disrupt the structural integrity of biofilms, enhancing the effectiveness of antibiotics such as vancomycin. These effects have been observed upon long exposure to hyperthermia (1-2 hours) and temperatures exceeding 50 °C [51]. In contrast, our reported NIRT effect achieves comparable results in a shorter duration and at lower temperatures.

Other researchers have explored the impacts of NIR employing a setup similar to ours (980 nm, 10 W, 148 J/cm²) on biofilms derived from clinical isolates of *S. aureus*[52].In contrast to our findings, they reported that NIRT did not influence biomass or bacterial viability significantly. However, a subtle disruptive effect on the biofilm structure was observed, aligning with the increased proportion of planktonic versus sessile/adherent bacteria following NIRT, as reported in our study.

In this context, our findings highlight that NIRT prompts the detachment of bacteria within the biofilm structure or strongly adhered to it, relocating them to the liquid phase of the culture. Simultaneously, NIRT induces a loss of biomass from the biofilm. This process exposes the released bacteria to both the PS and dissolved oxygen in the medium—critical components for bacterial inactivation during PDI. Additionally, it is established that heat enhances the permeability of prokaryotic cells. High temperatures have been demonstrated to alter staphylococcal biofilms by reducing their rigidity, thus favouring techniques used for their removal [51]. The observed detachment of bacteria from the biofilm, the observation of clefts after NIRT and the increased incorporation of PS post-NIRT, as evidenced in our results, align with these documented effects.Enhanced uptake of methylene blue was also documented after heating *E. coli* suspension at 46 °C, which was attributed to a modification in cell permeability [53].

The application of NIR facilitates rendering detached bacteria susceptible to treatment with antibiotics or PDI. In a study by Krespi et al. (2011), a successful combination of NIR therapy and shock waves was reported. This combination effectively disrupted *S. aureus* biofilms and eliminated the resulting planktonic bacteria using ciprofloxacin.

Furthermore, the successful combination of Infrared Radiation filtered by water (wIRA) with PDI utilizing different PSs, has been reported in various studies [54–56]. In contrast to the hyperthermia induced by wIR systems, in our work, NIRT achieves biofilm heating in a significantly shorter timeframe (21.5 °C increase in 30 sec). This rapid temperature rise, followed by cooling, induces the expansion and contraction of the biofilm matrix. This mechanical effect on the biofilm structure may contribute to the development of clefts, as observed in our results. These clefts would contribute to both the release of bacteria and the penetration of compounds from the liquid medium into the biofilm.

PDI is an alternative treatment option to combat infections, acting independently of complementary antimicrobial approaches, such as antibiotic therapy. However, complete eradication of microorganisms at the site of infection using PDI alone is extremely difficult [19].In addition, light-mediated technologies, however, might not necessarily supplant antibiotics but could serve to decrease the required doses or alleviate the pathogen load that needs elimination [57].

The findings presented in this study enable us to propose NIR irradiation as a supplementary treatment to PDI. This combined strategy of NIRT and PDI offers the advantage of reducing bacterial numbers using a lower concentration of PS compared to PDI as a standalone treatment, highlighting its potential as an effective and resource-efficient antibacterial approach.

## Materials and methods

### *S. aureus* biofilm cultures

We used the *S. aureus* RN6390 strain, a virulent derivative of the 8325 strain (Ingavale;Horsburgh). Cultures were grown in Trypticase soy agar (TSA)/broth (TSB) (DIFCO,USA) and stored in 20% glycerol TSB at −20 °C until use.

For the growth of biofilms, *S. aureus*strain was grown overnight in TSB and then diluted 1:1000 in 0.25% glucose-supplemented TSB. An aliquot of this cell suspension was inoculated into sterile (flat bottom) 24-well polystyrene microtiter plates (Corning, USA). After 24 hours of static incubation at 37°C, in aerobic conditions, biofilm formation was verified at the bottom of the well. Biofilms were carefully washed three times with sterile PBS and kept in 0.25% glucose-supplemented sterile PBS during experimental procedures adapted from [34].

### NIR laser device

We employed a LumiiaSonoBeam laser system with a 980 nm InGaAsP diode laser.The optical fibre has a core size of 600 µm and NA of 0.22, ensuring spot uniformity of ±15%. A lens (Thorlabs Inc., New Jersey, USA) was used to distribute the laser energy throughout the well diameter uniformly. This configuration resulted in an irradiation profile consisting of a cross-sectional area of 16 mm, which allowed homogeneous irradiation of the biofilms grown at the bottom of each well. The device emitted at 980 nm, and was employed in continuous or pulsed mode (2 Hz, 50 msec pulses). Light doses were adjusted by varying the output powers and/or the exposure times. Power measurement was conducted using a Coherent FieldMax II power meter with a Coherent PM150-50C head (Coherent Inc., Santa Clara, USA).

### Temperature measurement

The biofilm culture’s liquid medium (PBS) temperature was measured using a UNI-T Multimeter Model UT70A, recording changes in Celsius.

### Photosensitizer

TAPP was synthesized as described by Caminos et al [35]. Stock solutions of TAPP were prepared in dimethylformamide (DMF) and then diluted in water before use.

### Visible light source for PDI

Multiwell plates containing bacterial suspensions or biofilms were placed on glass and exposed to the light source from above and below using water filters and air-cooling to maintain temperatures below 20°C. Two ELH tungsten halogen GEQuartzline lamps with a reflector (500 W, General Electric Co., Cleveland, Oh, USA)were placed at a 25 cm distance from the sample, which provided a homogeneous total fluence rate of about 25.2 mW/cm^2^ on the surface of the sample measured with a FieldMaster power meter and a LM3 HTP sensor (Coherent Inc., Santa Clara, USA). The light dose was modified by switching the exposition time, which resulted in fluences between 22.5 and 180J/cm^2^.

### NIRT and PDI on biofilms

After growing the biofilms on the plates for 24 hours and performing the washes with PBS, as mentioned above, they were kept in PBS with 0.25% glucose to perform the different photo-assisted treatments. NIRT was carried out by irradiating the biofilm culture with the NIR laser device, under the conditions described above. To perform TAPP-PDI,TAPP was added to the biofilm cultures before NIR treatment. Afterwards, the biofilms were statically incubated in darkness at 37° C for one hour before being irradiated with the visible light source previously described.

After each treatment, the biofilms were scrapped off and the resulting suspensions were homogenised by vortex shaking. Bacterial viability (CFU/ml) was determined by serial dilutions of bacterial suspensions plated on TSA (colony counts were carried out after 24 hours of incubation at 37 °C).

In experiments involving the separation of the air-liquid phase (ALI) from the biofilms adhered to the well bottom post-NIRT, the liquid phase—comprising detached and weakly adhered bacteria—was promptly transferred to a fresh well. Concurrently, the remaining biofilm received a replenishment of sterile PBS containing 2.5 µM TAPP. Subsequently, PDI was conducted through individual irradiation with visible light for both phases. Following PDI, the quantification of CFU/ml was carried out in each phase, utilizing the previously described method.

### Biofilm TAPP binding/uptake assay

Biofilms were formed as explained above and TAPP was added to a final concentration of 2.5μM. Cultures were incubated for 1 hour at 37 °C and subsequently, scrapped biofilmsuspensions were centrifuged for 10 min at 13,000 g. The supernatants were removed and the remaining pellets were washed twice with PBS. The washed pellets were then mixed with DMSO at room temperature and sonicated. The mixtures were centrifuged for 10 min at 13,000 g and the DMSO supernatants were removed for fluorescence analysis. Fluorescence was determined at the excitation wavelength set to 419 nm and the emission wavelength set to 660 nm after analysis of the emission spectrum (PerkinElmer LS55 fluorimeter, UK). The concentration of TAPP was determined through the use of standard curves of fluorescence versus the concentration of TAPP dissolved in DMSO. This value was then divided by the CFUs of each culture.

### Crystal violet assay

Crystal violet staining was used to quantify biofilm biomass. Briefly, after performing the treatments with the NIR laser, the supernatants of the cultures were discarded and three washes were performed with sterile PBS. Then, the attached cells were fixed with methanol for 15 min, stained with crystal violet solution (0.1%) and incubated statically for 20 min at room temperature. The dye solution was removed and four washes were performed with distilled water. After air drying for 30 minutes at room temperature, acetic acid solution (30%) was added. Finally, the absorbance was measured at 595 nm.

### Scanning electron microscopy (SEM)

*S. aureus* biofilms were cultured on coverslips using the previously outlined procedure. Following each experimental treatment, samples underwent fixation with 4% glutaraldehyde in 0.1 M cacodylate buffer at pH 7.2, for 2 hours at 4°C. Subsequently, they underwent a series of ethanol incubations at escalating concentrations (25%, 50%, 75%, and 96%) for 10 minutes each, followed by air-drying at room temperature. Ultimately, the specimens were coated with Au-Pd using a Thermo VG Scientific SC 7620 Sputter Coater and subjected to examination via SEM utilizing a Philips model XL30 TMP (Eindhoven, The Netherlands).

### Biofilm microscopy analysis of viability

Biofilms were developed on sterile coverslips placed in multi-well plates and treated with NIRT or PDI as indicated previously. To determine the viability of bacteria within the biofilms after the different treatments, the LIVE/DEAD BacLight Bacterial Viability Kit (Molecular Probes, Eugine, OR) was used as previously described (Mamone et al 2016). The biofilms were examined by confocal scanning laser microscopy(CSLM) employing a Zeiss LSM 510 Meta confocal microscope. Optical sections of 0.56 μmwere collected, and for each sample, images from three randomly selected positions were acquired and processed using the ImageJ software (Wayne Rasband, National Institutes of Health).

### Monitoring of mice skin temperature during *in vivo* NIRT

Male CF-1 mice, 8 weeks old, weighing 20–25 g, were used. They were provided with food(Molinos Rio de la Plata) and water ad libitum. Mice were anaesthetized by intraperitoneal injection of 70 mg/kg ketamine hydrochloride and 6 mg/kg xylazine. Then, the skin of the left flanks was shaved and treated with a brief spurt (1 second) of cryogen spray (Nassau, Buenos Aires, Argentina) to impart a cooling and anaesthetic effect on the epidermis and the surrounding tissues.Finally, irradiation was carried out with a LumiiaSonoBeam laser for 30 sec at 7.5 W/cm^2^, continuous mode. The spot size comprised a skin area of 1 cm^2^. The temperature at the skin surface was followed by using a UNI-T Multimeter Model UT70A placed over the skin.

Animal protocols were approved by the Argentinean Committee (CICUAL, School of Medicine, University of Buenos Aires, Res CS 4081/04). The reporting of *in vivo* experiments follows the recommendations in the ARRIVE guidelines.

### Statistics

Values were expressed as mean ± standard deviations of the mean. One-way and two-way Analysis of Variance (ANOVA) with post hoc comparisons (Tukey) were used to determine statistical significance. To assess the interaction between the factors PDI and NIRT, data analysis was conducted using a two-way ANOVA, evaluating both the effects of each factor separately and the interaction factor. The corresponding P-values for the different tests are reported in the figures. In all cases, statistical tests were carried out with the GraphPad Software (version 5.0; GraphPad Prism). P values: * ˂0.05, ** ˂0.01, *** ˂0.001 and **** ˂0.0001were considered significant.

## Abbreviations

ALI: air-liquid interface
CFUs: colony forming units
CSLM: confocal scanning laser microscopy
PBS: Phosphate-buffered saline
NIRT: Near Infra-red Treatment
NIRTc: NIRT in continuous mode
NIRTf: fractionated NIRT
NIRTp: NIRT in pulsed mode
PDI: Photodynamic Inactivation
PS: photosensitizer, photosensitizers
SEM: scanning electronic microscopy
TAPP: 5,10,15,20-tetrakis[4-(3-*N*,*N-*)dimethylaminopropoxy)phenyl]porphyrin
TAPP-PDI: TAPP mediated photodynamic inactivation
TSA: Trypticase Soy agar
TSB: Trypticase Soy Broth.

## ACKNOWLEDGEMENTS

LM, GDV, FB, ED and AC are members of the Research Career of CONICET, Argentina. RT is a fellow at CONICET. AC thanks to ANPCyT (PICT 0727/18) and CONICET (PIP2014 number 11220130100237CO). EDthanks to ANPCYT (PICT 02391/19) and CONICET(PIP2021 number 11220200101208CO).We are grateful to Miss Vanina Ripoll for her technical support and to Lumiia S.A. for the provision of the 980 nm laser and technical support.

